# HAK-actin, U-ExM-compatible probe to image the actin cytoskeleton

**DOI:** 10.1101/2025.08.26.672318

**Authors:** Olivier Mercey, Luc Reymond, Florent Lemaître, Isabelle Mean, Marine H. Laporte, Marine Olivetta, Karin Sadoul, Omaya Dudin, Virginie Hamel, Paul Guichard

## Abstract

Expansion microscopy (ExM) enables super-resolution imaging by physically enlarging biological samples. While ExM has been successfully applied to study the intracellular microtubule cytoskeleton, reliable probes for visualizing actin fibers remain limited. Here, we present HAK-actin, an engineered actin probe compatible with post-labeling Ultrastructure Expansion Microscopy (U-ExM). We show that HAK-actin delivers robust and uniform actin staining across diverse systems, including human cells, microbial eukaryotes, and mouse retinal tissue. This tool provides a simple, versatile, and reproducible solution for actin cytoskeleton visualization, addressing a critical need in cell biology.

## Main Text

Expansion microscopy revolutionized the super-resolution field, giving access to the nanoscale molecular organization of the cell by magnifying the biological specimen itself^1–4^. The microtubule network has been largely characterized by expansion microscopy^3–7^, however, very few applications have targeted actin filaments, yet widely studied across biology fields. To date, actin visualization using expansion microscopy has been achieved using antibodies directed against either endogenous actin^3,8^ or genetically encoded actin probes, such as IntAct^9^ or Lifeact ^3^. However, their applicability is limited: antibody-based methods depend on the availability of species-specific reagents, while genetically encoded probes require efficient transfection or genetic manipulation. More recently, an alternative strategy has been developed to specifically visualize F-actin using phalloidin conjugated to trifunctional linkers (such as TRITON), which can be grafted onto hydrogels for ExM^10–12^. This method offers the advantage of leveraging phalloidin’s high affinity for F-actin, enabling visualization across multiple species. However, these designed linkers utilize the succinimidyl ester of 6-((acryloyl)amino)hexanoic acid (AcX) as an anchoring agent, which is not compatible with all ExM protocols. Additionally, they rely on a pre-expansion labeling strategy that constrains fluorophore selection^13^, leads to volumetric signal dilution after expansion^14^, and ultimately limits the achievable resolution^5,15^. Furthermore, this pre-labeling approach is incompatible with some protist species protected by protective structures such as cell walls, which often hinder labeling permeability^16,17^. These limitations highlight the need for more versatile and universally compatible linker chemistries to enhance the applicability and performance of ExM-based imaging approaches.

Here, we aimed to develop actin probes that are compatible with post-labeling expansion, thereby overcoming fluorochrome limitations and minimizing signal dilution. To achieve this, we designed a probe suited for the ultrastructure expansion microscopy (U-ExM) protocol^18^, which is adapted from the Magnified Analysis of the Proteome (MAP) method^19^. MAP preserves the entire proteome, enabling post-expansion labeling. U-ExM is particularly adapted for studying the actin cytoskeleton, as it maintains the molecular architecture of subcellular structures. U-ExM is also compatible with cryo-fixation (Cryo-ExM), an extension of the method that avoids artifacts due to chemical fixation, thereby optimizing cellular architecture preservation^3^. Moreover, U-ExM does not require specific anchoring molecules, as the acrylamide within the swellable polymer is directly cross-linked to primary amines of the entire proteome using simple formaldehyde treatment. Based on this property, we hypothesized that incorporating lysine residues, containing primary amines, into the well-characterized F-actin probes Jasplakinolide (JASP) or Phalloidin (PHALLO) would facilitate their retention within the hydrogel, as previously proposed^1,20^.

For this purpose, we first synthetized two JASP-based probes, both carrying an Alexa488 fluorochrome, known to withstand expansion process^21^, one probe incorporated four lysine residues (JASP-K4-488), while the other did not (JASP-488) (**Figure 1A**). Both probes were assessed in the osteosarcoma cell line U2OS, where actin cytoskeleton staining is easily assessed. After paraformaldehyde (PFA) fixation and cell permeabilization, actin probes were added for an hour, followed by either direct immunofluorescence or embedding in the swellable hydrogel for U-ExM (**Figure 1B**). After successful validation of these probes in regular immunofluorescence (**Supplementary Figure 1A**), we analyzed their retention after expansion. While JASP-488 produced no signal compared to the anti β-actin antibody control, the addition of lysine residues enabled JASP-K4-488 to effectively anchor to the hydrogel, as demonstrated by an increase in fluorescence intensity (**Figure 1C, D, G**). Despite this improvement, the signal appeared weak compared to background, likely due to the volumetric dilution of the fluorochrome within the expanded hydrogel.

**Figure 1.**
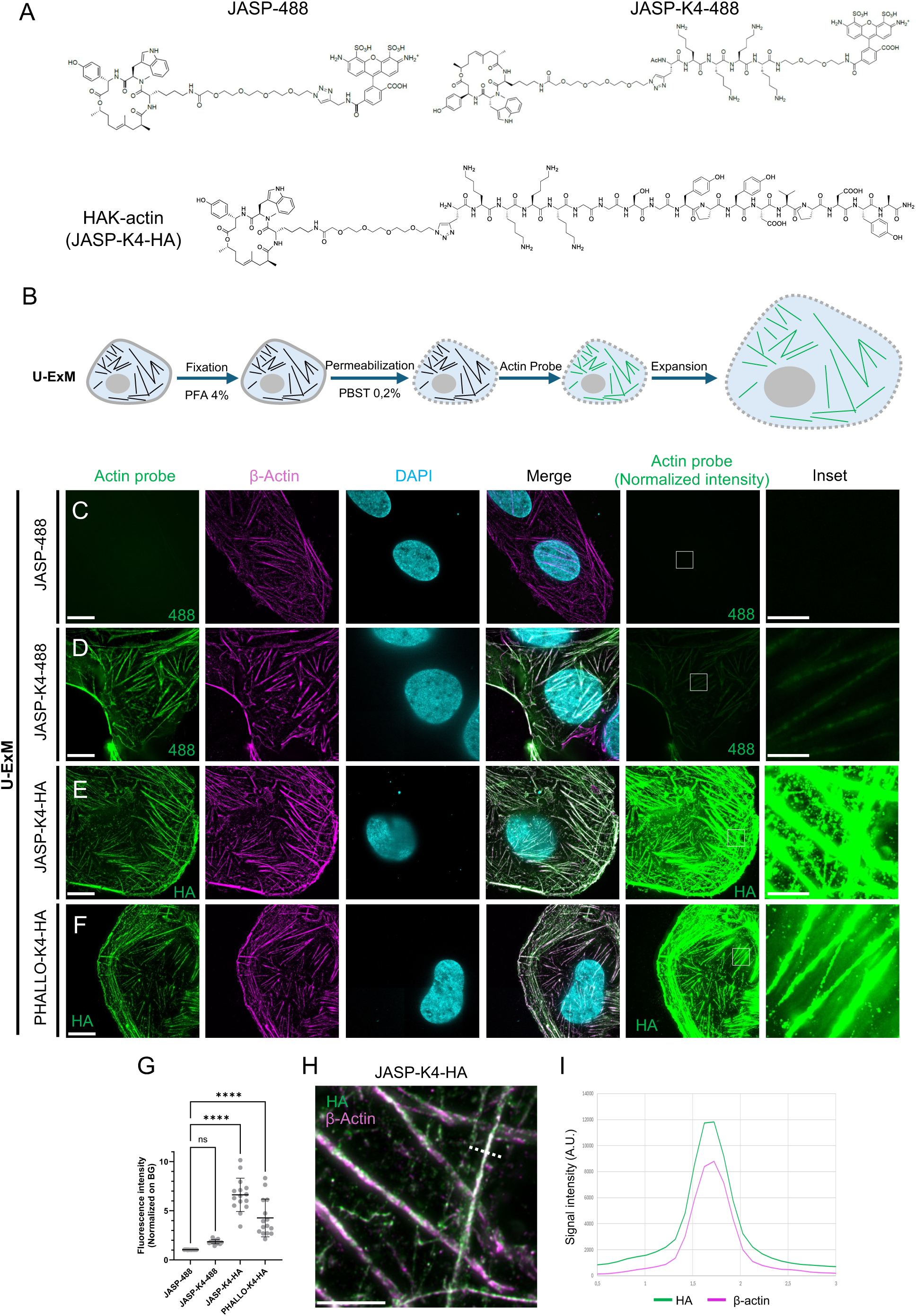
HAK probe efficiencies in expanded U2OS cells. **A**. Chemical structures of three actin probes designed for U-ExM and mainly used in this study. All of them are composed of the Jasplakinolide (JASP) molecule coupled to a PEG-moiety and a propargylglycine. On JASP-488, an Alexa 488 fluorophore is directly coupled to JASP. Both JASP-K4-488 or JASP-K4-HA (HAK-actin) have been designed with four lysine residues and an Alexa 488 or an HA-tag, respectively. **B**. Workflow of HAK-actin probe use in U-ExM. The probe is added for 1h after cell fixation (PFA 4% and permeabilization with PBS Tween 0,2%), and cells are then processed for expansion. **C-F**. Representative widefield images of PFA-fixed and expanded U2OS cells treated with different actin probes. Actin probe staining (probe directly coupled to the Alexa 488 fluorophores or revealed through immunostaining post expansion using anti-HA antibodies) is shown in green, anti-Beta-actin staining is shown in magenta and nuclei in cyan (DAPI). Scale bars: 10 µm. The right panel shows the HAK-probe signals normalized among all condition, and an inset is depicted just on the right. Scale bar: 2µm. **G**. Quantification of the fluoresence intensity normalized against background of actin probes in the different conditions analyzed in C-F (n=15 cells with 3 independent experiments per conditions). **H**. Inset of the JASP-K4-HA (HAK-actin) to highlight the overlap between the actin probe signal and beta actin staining. Scale bar: 2µm. **I**. Plot profile resulting from the dashed line in H.

To address this major limitation, frequently reported in various expansion microscopy experiments^13^, we added a post-fixation step using PFA and Glutaraldehyde (GA). However, while this approach improved the signal intensity, the overall enhancement remained modest (**Supplementary Figures 1B, C**). To further improve signal intensity, we designed and synthetized a novel actin probe for post-expansion labeling by incorporating a Hemagglutinin (HA) tag (JASP-K4-HA) (**Figure 1A**). The HA tag is widely used in cell biology as an epitope tag for protein detection and purification via immune recognition with commercial antibodies. It consists of a short peptide sequence (YPYDVPDYA) derived from the HA protein of the influenza virus. As expected for this post-labeling strategy, JASP-K4-HA enabled clear visualization of actin fibers and produced strong signal amplification (**Figure 1E**). Moreover, this probe offers flexibility in fluorophore selection, as it can be combined with conventional secondary antibodies that can be conjugated to a wide range of fluorochromes. We refer to this new design as HAK, reflecting the incorporation of both an HA tag and lysine (K) residues.

To evaluate the signal enhancement achieved through post-expansion immunostaining, we measured the fluorescence intensities of the different probes in expanded U2OS cells. Our measurements revealed a four-fold increase in signal intensity for JASP-K4-HA compared to JASP-K4-488, demonstrating the enhanced signal provided by the HA tag in the HAK probe design (**Figure 1E, G**). To test the versatility of this approach, we next designed a Phalloidin (PHALLO)-based probe, following the same strategy (PHALLO-K4-HA) (**Supplementary Figure 2** and **Figure 1F, G**). While we noticed that PHALLO-K4-HA displayed a lower signal (∼1.5 fold decrease) compared to JASP-K4-HA, we measured a two-fold increase in signal intensity compared to a JASP-K4-488 probe (**Figure 1F, G**). Importantly, the specificity of both HAK-probes was confirmed by their colocalization with the immuno-labeled β-actin (**Figure 1H-J**).

Encouraged by the efficiency of the HAK probe design, we next explored whether alternative types of retention molecules could further enhance probe anchoring. We tested different amino acids bearing a side chain amino group: Lysine (K4, original design), Diaminopropionic acid (DAP), Ornitine (ORN) and 4-aminophenylalanine (4AF) (**Figure 2A** and **Supplementary Figure 2**). Signal intensity measurements, used as an indicator of anchoring efficiency, revealed that K4 and 4AF residues provided the highest retention within the hydrogel (**Figure 2A, B**). Therefore, the best HAK-probe design chosen for this study, JASP-K4-HA, was dubbed HAK-actin. It includes the Jasplakinolide (JASP) actin ligand, four lysine residues, and an HA tag (**Figure 1A**).

**Figure 2.**
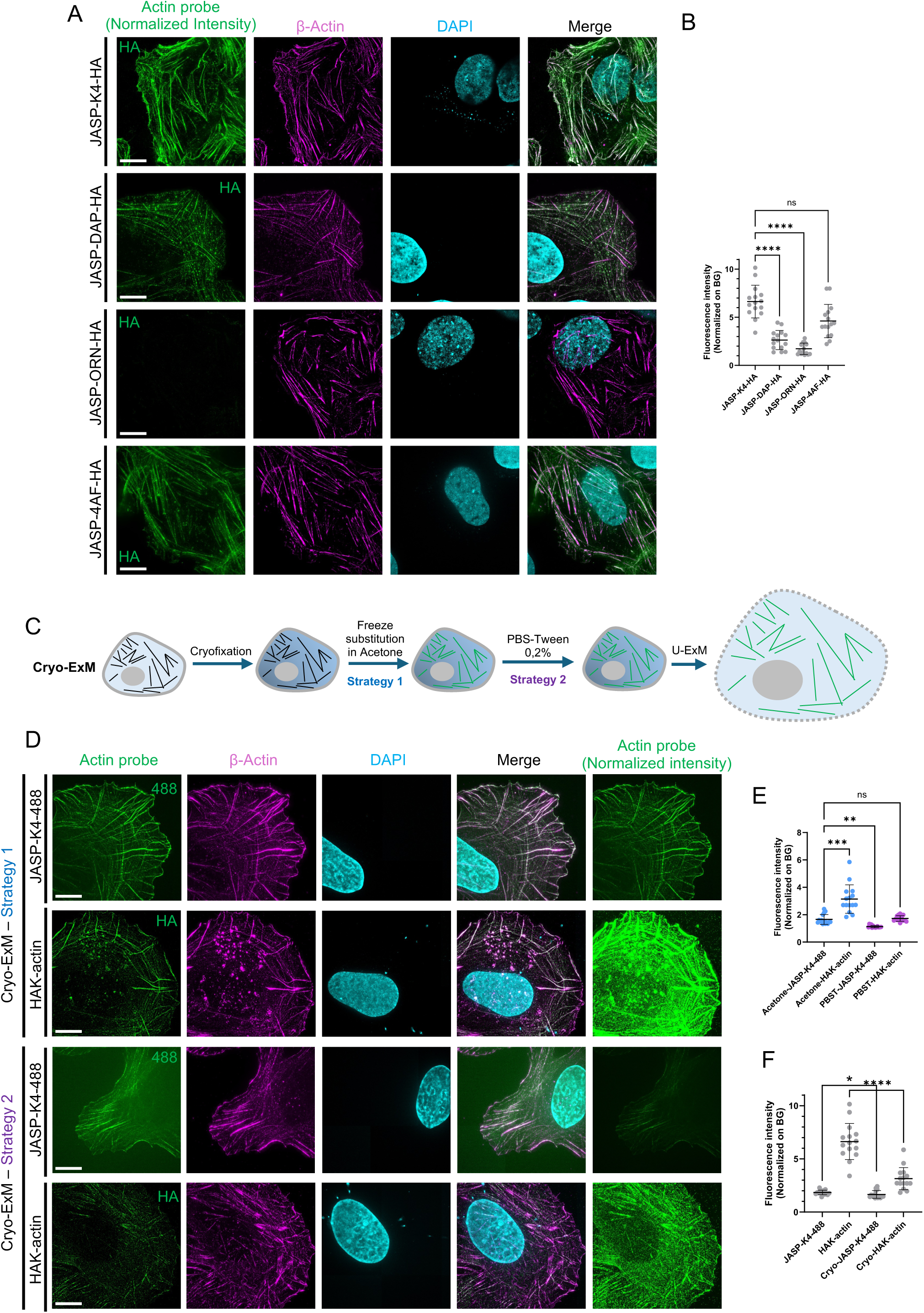
Optimization of HAK probes with different linkers and fixation conditions. **A**. Representative widefield images of expanded U2OS cells treated with several actin probes bearing different anchoring linkers: Lysines (K4), Diaminopropionic acid (DAP), Ornitine (ORN) et 4-aminophenylalanine (4AF). Actin probes (normalized across images) are shown in green, Beta-actin antibodies in magenta and DAPI in cyan. Scale bars: 10 µm. **B**. Quantification of the fluorescence intensities normalized to background for the different probes (n=15 cells; N=3 independent experiments per conditions). **C**. Workflow of HAK-actin use in cryo-ExM. The probe is added either during the freeze substitution process in acetone (strategy 1), or in PBS-Tween 0,2% just before the expansion microscopy process (strategy 2). **D**. Representative widefield images of cryo-fixed and expanded U2OS cells treated with either JASP-K4-488 or HAK-actin during the freeze substitution step (strategy 1) or just before the expansion process (strategy 2). Actin probes are shown in green (normalized across images on the right), Beta-actin antibodies in magenta and DAPI in cyan. **E**. Quantification of the signal intensities (normalized on background) in the different cryo-conditions (n=15 cells; N=3 independent experiments per conditions). **F**. Quantification of the signal intensities (normalized on background) observed with cryo-fixation conditions or with PFA fixation (n=15 cells; N=3 independent experiments per conditions). Note that some values are the same as in Figure 1G and Figure 2E for direct comparison.

Since chemical fixation can introduce artifacts in cell architecture^22,23^, we next tested the compatibility of these actin probes with Cryo-ExM, which better preserves cellular ultrastructure through cryo-fixation and freeze substitution^3^ (**Figure 2C-F**). We incorporated actin probes JASP-K4-488 or HAK-actin into either the acetone during the freeze substitution process (strategy 1) or in PBS-Tween 0,2% just before the expansion microscopy process (strategy 2) (**Figure 2C)**. Both strategies successfully lead to the detection of an actin signal, but incorporation during freeze substitution appears to be significantly more efficient (**Figure 2D, E**). Consistently, we also observed an amplification of the signal using HAK-actin compared to JASP-K4-488 (**Figure 2E**), further highlighting the efficiency of the post-labeling strategy of the HAK-probe regardless of the fixation conditions. We thus concluded that the HAK probes are compatible with cryo-ExM, although the signal was slightly lower than under PFA fixation (**Figure 2F**). This compatibility represents a significant advantage for imaging cellular architecture, as Cryo-ExM can also be used to visualize membranes by incorporating BODIPY^24^. To assess this, we tested the approach and successfully obtained images where both membrane (stained with mCLING^25,26^) and actin cytoskeleton were clearly labeled (**Supplementary Figure 3**). We further applied this approach to image the cleavage furrow, a transient, actin-rich structure that participates in daughter cell separation during cytokinesis (**Figure 3A**). We next evaluated the compatibility of HAK-actin with the iterative iU-ExM protocol, which achieves up to 16-fold expansion of biological specimens^4^. HAK-actin successfully labeled actin filaments in U2OS cells expanded using iU-ExM, demonstrating its suitability for high-resolution visualization under these conditions (**Figure 3B**).

**Figure 3.**
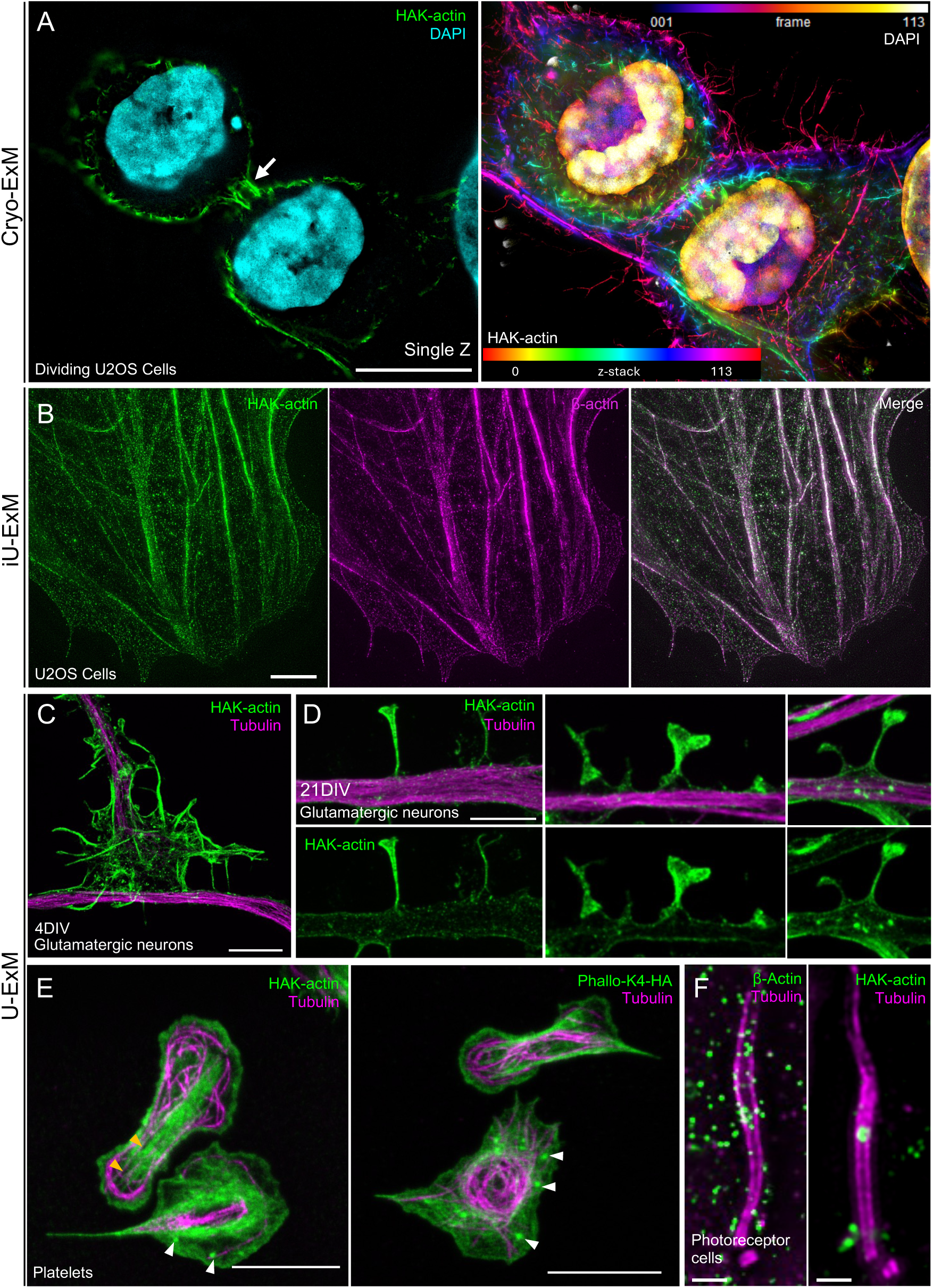
HAK-actin in several mammalian cells and tissue models. **A.** Widefield image of PFA-fixed expanded U2OS cells labeled with HAK-actin and DAPI, highlighting the actin enrichment at the level of the cleavage furrow. Left panel is a full Z-Projection of the cells where HAK-actin is color-coded with “spectrum” and DAPI is color-coded with “fire” for z positions, whereas the right panel is a single z image plane highlighting the intense actin staining (Green) at the cleavage furrow (white arrow). Scale bar: 10µm **B.** A widefield image of cryo-fixed U2OS cells processed for iU-ExM and stained for HAK-actin (Green) and beta actin (magenta). Scale bar: 2 µm. **C**. Confocal images of expanded immature (4DIV) glutamatergic neurons stained with the HAK-actin (green) and tubulin (magenta) revealing the actin network in growth cones of immature neurons. Scale bar: 5 µm. **D**. Confocal images of expanded mature (21DIV) glutamatergic neurons stained with the HAK-actin (green) and tubulin (magenta) showing dendritic spine diversity in mature neurons (21 DIV). Scale bar: 2,5 µm. **E**. Widefield images of expanded human activated platelets stained with tubulin in magenta and HAK-actin (left) or Phallo-K4-HA (right) in green. Stress fibers are highlighted with orange arrowheads and actin nodules with white arrowheads. Scale bar: 5 µm. **F**. Widefield images of expanded murine connecting cilium from photoreceptor cells stained with tubulin in magenta and beta actin (left) or HAK-actin (right) in green. Scale bar: 500 nm.

To further assess the versatility of HAK-actin, we tested its performance in various mammalian cells and tissue models. We first investigated human glutamatergic neurons at different stages of differentiation, where actin network plays a critical role in organizing distinct subcellular compartments (**Figure 3C, D**). In immature neurons (4 DIV, **Figure 3C**), the actin network is particularly important for the structural organization of growth cones, which facilitate connections with their target neurons during development^27^. Application of HAK-actin on expanded neurons enabled visualization of the canonical ultrastructure of growth cones, revealing peripheral actin fibers, lamellipodia, and filopodia surrounding the microtubule network, as well as their interactions with neighboring cells (**Figure 3C**). At mature stage (21DIV, **Figure 3D**), glutamatergic neurons start to establish dendritic spines, specific structures crucial for synaptic transmission. The actin cytoskeleton provides structural integrity to these spines, dictating their characteristic morphologies. Here again, HAK-actin labeling on expanded neurons allowed for the identification of dendritic spines exhibiting different morphological subtypes, including thin, mushroom, and cup-shaped spines, as previously reported in the literature^28^.

We next imaged activated human platelets, which undergo massive actin reorganization from actin nodules, i.e. a podosome-like organization^29^ to actin stress fibers crossing the platelet longitudinally^30^. While in non-expanded platelets, a precise actin organization was not visible (**Supplementary Figure 4A)**, the use of HAK-probes, HAK-actin or PHALLO-K4-HA, on expanded platelets clearly revealed two typical examples of actin organizations, stress fibers and actin nodules (**Figure 3E**, **Supplementary Figure 4A**, orange and white arrowheads, respectively)^30^.

To extend these findings to more complex tissues, we next examined actin labeling in mouse retina that we previously studied using U-ExM^31–33^. Recent studies suggest that actin accumulates above the connecting cilium of photoreceptor cells and participates in the membrane disc formation of the outer segment cilium, crucial for visual acuity^34^. We compared the actin signal obtained using an anti β-actin antibody to the signal obtained with HAK-actin, followed by post-expansion immunolabeling with anti-HA antibodies (**Figure 3F**). Both strategies exhibited an actin signal at the level of the bulge region^33^, where microtubules enlarge and where membrane discs are formed. Notably, HAK-actin yielded a clearer signal with reduced background, validating its utility for actin imaging in expanded tissue samples.

Finally, to determine whether HAK-actin can be used as a broad-spectrum U-ExM actin markers, we tested its efficiency in microbial eukaryotes. Microbial eukaryotes display diverse cytoskeletal features due to their flexible life cycles and are often impermeable to antibody labeling due to protective layers like cell walls. However, recent work showed that U-ExM enhances antibody accessibility for immunostaining and improves resolution as previously shown for microtubules^17,35–37^. Genetic tagging provides an alternative approach, but many species remain genetically intractable, and antibodies recognizing actin are often unavailable or species-specific. We first focused on the ichthyosporean *S. arctica*, one of the closest living relatives of animals, in which phalloidin-based staining has revealed a filamentous actin network decorating furrow ingressions during cellularization, similar to early *Drosophila* embryogenesis^38–40^. We compared unexpanded and U-ExM-expanded *S. arctica* cells labeled with JASP-488, JASP-K4-488, or HAK-actin, followed by conventional immunofluorescence or U-ExM (**Figure 4A**). We found that JASP-488 stains unexpanded cells but fails to label actin in expanded samples, as expected, since this probe is not retained within the hydrogel (**Figure 4A, B**). JASP-K4-488, while most likely retained in the hydrogel, was insufficient to improve staining efficiency in expanded cells, likely due to a loss of fluorophore intensity under U-ExM conditions. By contrast, HAK-actin, detected with anti-HA antibodies, was not visible in unexpanded cells but efficiently labeled expanded *S. arctica* cells, clearly highlighting actin at furrow ingressions during cellularization (**Figure 4A-C**). Thus, HAK-actin, via post-expansion immunolabeling, enables visualization of actin structures in this species that conventional anti-actin antibodies could not reveal, even after expansion (**Figure 4B, C**).

**Figure 4.**
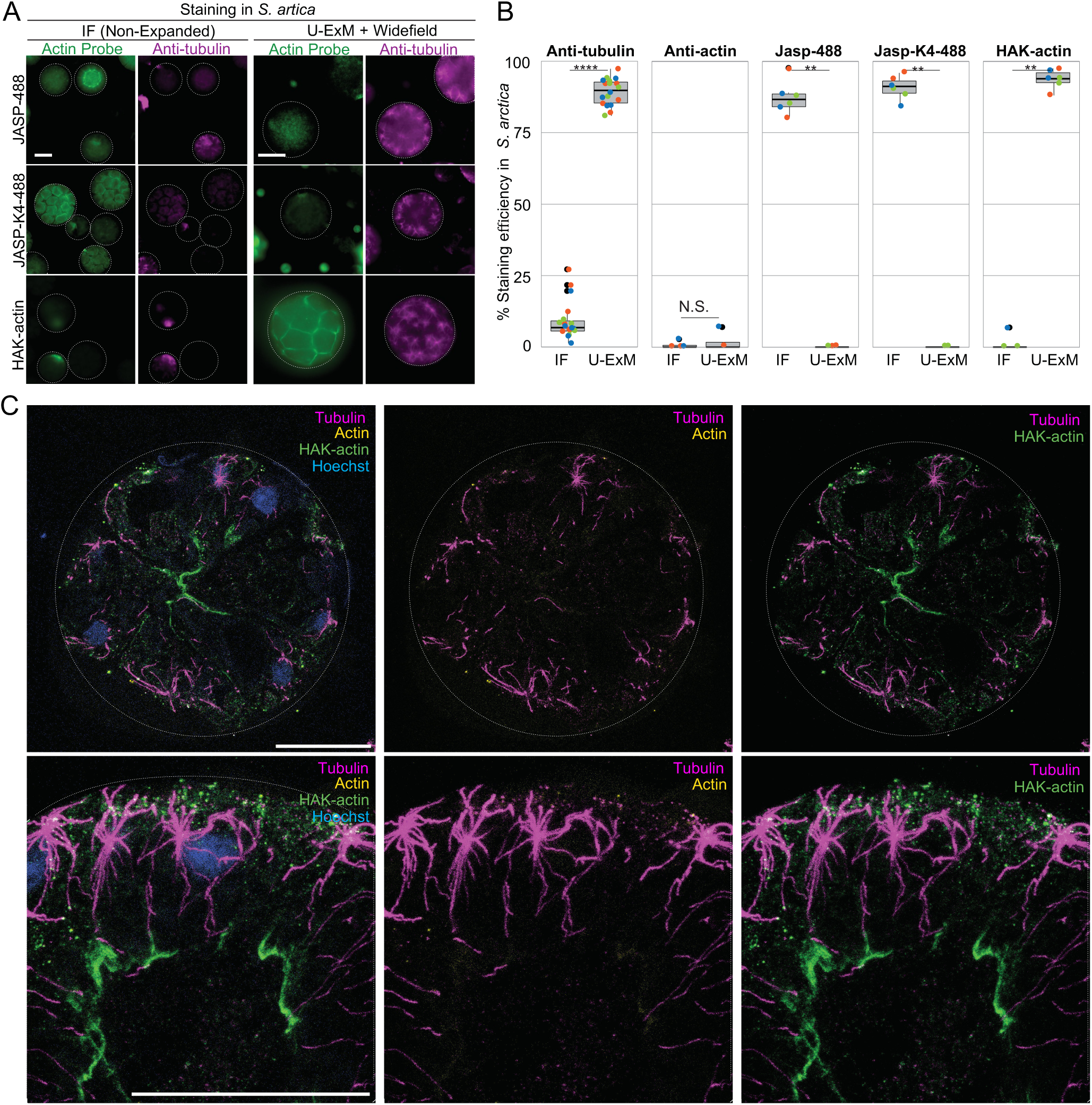
Unveiling actin network in Microbial Eukaryotes. **A**. *Sphaeroforma arctica* cells labeled for actin (green) using JASP-488, JASP-K4-488 and HAK-actin and for tubulin (magenta) imaged as non-expanded samples, or after U-ExM, using widefield microscopy. Note that U-ExM enables both antibody, and probe-based staining in this cell-walled species, revealing its actin and microtubule networks. Scale bar: 10 µm. **B**. Boxplot showing the efficiency of antibody, and probe-based staining for actin and tubulin in *S. arctica* under non-expanded (immunofluorescence, IF) versus expanded (U-ExM) conditions. While U-ExM facilitates tubulin immunostaining, actin antibodies do not stain effectively, whereas HAK-actin in combination with an anti-HA antibody successfully label filamentous actin (n>30 cells per replicate, 3 independent replicates were performed).**C**. U-ExM samples of *S. arctica* undergoing actomyosin–dependent cellularization, stained with the HAK-actin in combination with an anti-HA antibody (green), anti-tubulin (magenta), anti-actin (yellow), and Hoechst for DNA. The actin furrows typical of cellularization are clearly visible using the probe but difficult to detect with actin antibodies. Scale bar: 10 µm.

We next assessed the impact of a post-fixation step using 4% formaldehyde and 0.025% glutaraldehyde after incubation of pre-cellularizing *S. arctica* cells, where actin localizes as small cortical actin nodes^39,40^ with JASP-K4-488 or HAK-actin, prior to U-ExM (**Supplementary Figure 4B, C**). Our results show that this crosslinking step enhances actin labeling with HAK-actin but not with JASP-K4-488, suggesting that both post-fixation step and HA-tagged actin probe are required for optimal actin detection in expanded cells.

We further tested other microbial eukaryotes including two additional ichthyosporeans, *Creolimax fragrantissima*^41^ and *Chromosphaera perkinsii*^42^, as well as the green alga *Volvox tertius*^43^, and the amoebozoan *Amoeba proteus*^44^, (**Supplementary Figure 4D**). Consistent with previous findings, we identified filamentous actin structures in all tested species (**Supplementary Figure 4D**). In *C. fragrantissima* and *C. perkinsii*, actin localized at cell boundaries, whereas in *V. tertius*, we observed perinuclear actin rings and actin co-localizing with the flagellum^45^. In *A. proteus*, despite the absence of tubulin labeling, likely due to a specific cell stage lacking microtubules, we detected an extensive actin network (**Supplementary Figure 4D**)^46^. Additionally, we attempted to label actin in the ciliate *Tetrahymena pyriformis*^47^ and the euglenid *Euglena gracilis*^48^, but observed no specific staining (**Supplementary Figure 4D**). This result aligns with previous reports that phalloidin-based actin labelling fails in both species^49–53^, suggesting a lack of filamentous actin in these lineages.

Together, these results suggest that HAK-actin serves as a broad-spectrum marker for visualizing filamentous actin in U-ExM-expanded samples across diverse microbial eukaryotes.

## Discussion

In this study, we developed and validated HAK-actin, a novel probe optimized for actin labeling in Ultrastructure Expansion Microscopy (U-ExM). By engineering the well-characterized actin ligand Jasplakinolide with additional lysine residues and an HA tag, we achieved strong probe retention and robust fluorescence signal in expanded samples through post-expansion labeling with conventional antibodies. This approach enables both signal amplification and greater flexibility in fluorophore selection. Our results show that HAK-actin overcomes key limitations of existing ExM-compatible actin markers, including fluorophore signal dilution and the restrictions associated with pre-expansion labeling. Importantly, our approach is compatible with U-ExM, cryo-ExM, and iU-ExM. We show that HAK-actin reliably labels actin filaments in both chemically fixed and cryo-fixed samples. Furthermore, we demonstrate the versatility of this probe across diverse biological systems, ranging from mammalian cells and tissues to microbial eukaryotes.

Beyond their utility for structural imaging, HAK-actin opens new possibilities for investigating actin organization in expanded samples. The HA tag enables selective post-expansion immunostaining, offering the potential for multiplexed imaging approaches that combine actin visualization with other cellular components. Additionally, the ability to pair HAK-actin with cryo-ExM allows for more accurate preservation of cytoskeletal architecture, which is particularly valuable for studying highly dynamic processes such as cell division, migration, and intracellular trafficking. Owing to its straightforward cross-linking and post-expansion labeling strategy, which is compatible with a broad range of expansion microscopy protocols, this probe holds strong potential for adaptation across diverse ExM methodologies.

In conclusion, the development of HAK-actin represents a significant step forward in the application of expansion microscopy for actin imaging. Their versatility, compatibility with multiple U-ExM protocols, and broad-spectrum applicability across different biological models establish them as a valuable tool for high-resolution cytoskeletal studies.

## Acknowledgments

We thank M. Delous, F. Lecaignard (CRNL), S. Blondel and A. Ruiz (GenCyTi, CRNL) for the culture iPSCs-derived glutamatergic neurons. We thank Umut Batman for initial experiments. We acknowledge the contribution of SFR Santé Lyon-Est (UAR3453 CNRS, US7 Inserm, UCBL) facility: CIQLE (a LyMIC member) for the acquisition of neuron images. We also thank E. Paulin for the preliminary results obtained with cryo-expansion on U2OS. This work was supported by the Swiss State Secretariat for Education, Research and Innovation (SERI) under contract number MB22.00075 (PG), the Swiss National Foundation (SNSF) 310030_205087 attributed to P.G. and V.H. and the Pro Visu, Gelbert and TANDEM ISREC Foundations attributed to V.H. M.O. and O.D., were supported by a Swiss National Science Foundation Starting Grant (TMSGI3_218007).

## Author Contributions

L.R, P.G. and V.H designed the HAK probes, which were synthetized by L.R.. F.L, I.M and O.M designed and performed experiments on U2OS cells. K.S and M.H.L performed the experiments on platelets and neurons. O.D and M.O performed experiments on protists. OM performed experiments on retina. P.G and V.H supervised the study. O.M assembled the figures; O.M, V.H and P.G. wrote the manuscript with inputs from all co-authors.

## Competing financial interests

LR, PG, OD, and VH are inventors of a patent application (EP25184605.1) assigned to University of Geneva covering HAK probes. L.R. owns shares of Spirochrome A.G., commercializing HAK-actin probe. The other authors declare no Competing financial interests.

---

**Supplementary Figure 1.**
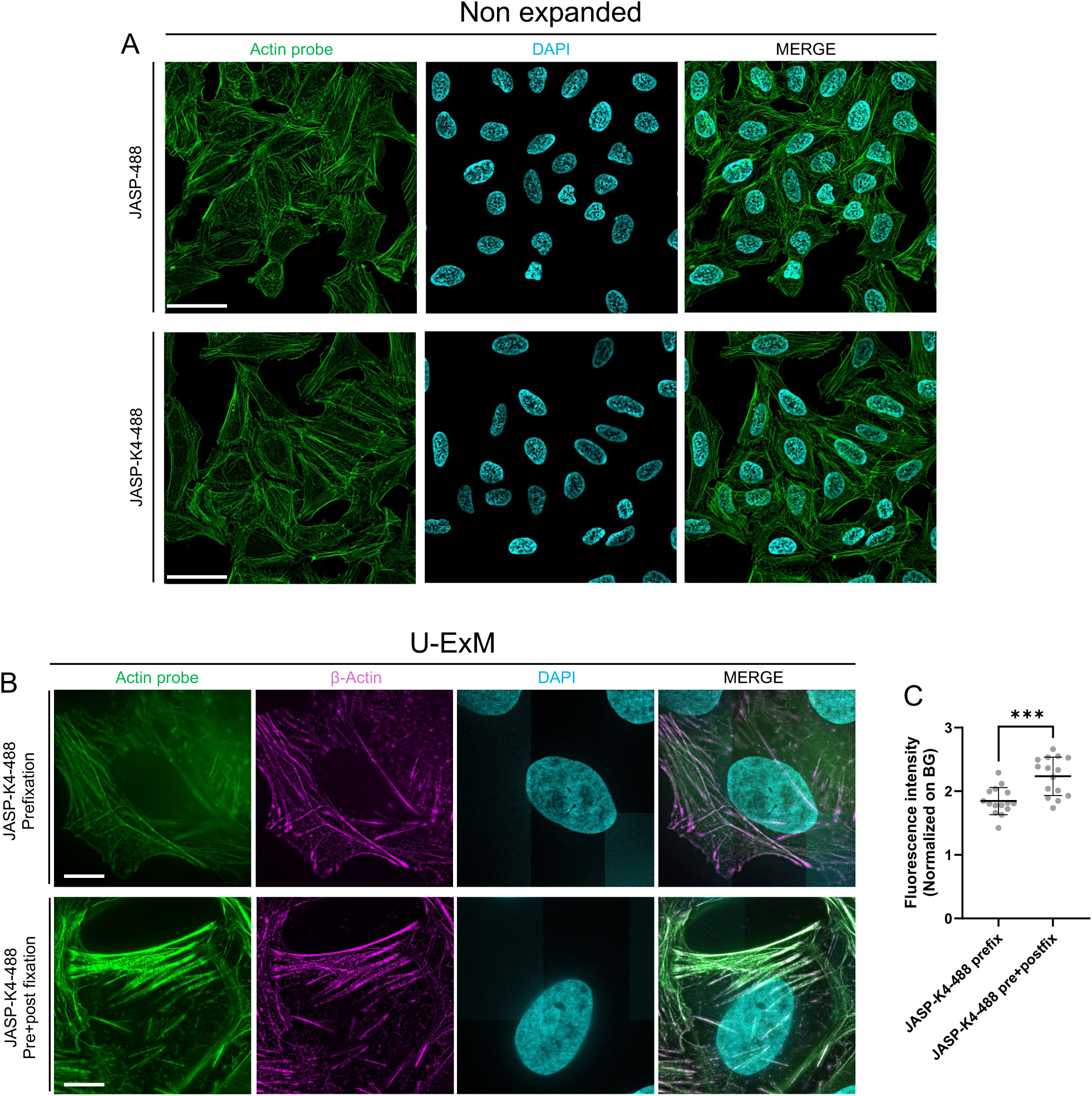
Additional tests to evaluate the actin probes under different conditions. **A**. Immunofluorescence images of non-expanded U2OS cells treated with either JASP-488 or JASP-K4-488 (green) together with DAPI (cyan). Scale bar: 50 µm. **B**. Widefield images of expanded U2OS treated with JASP-K4-488 (green) with or without post-fixation step (PFA 4%, Glutaraldehyde 0,0125%). Scale bar: 10 µm **C**. Signal intensity measurement of JASP-K4-488 assessing the effect of post fixation (n=15 cells; N=3 independent experiments per conditions).

**Supplementary Figure 2.**
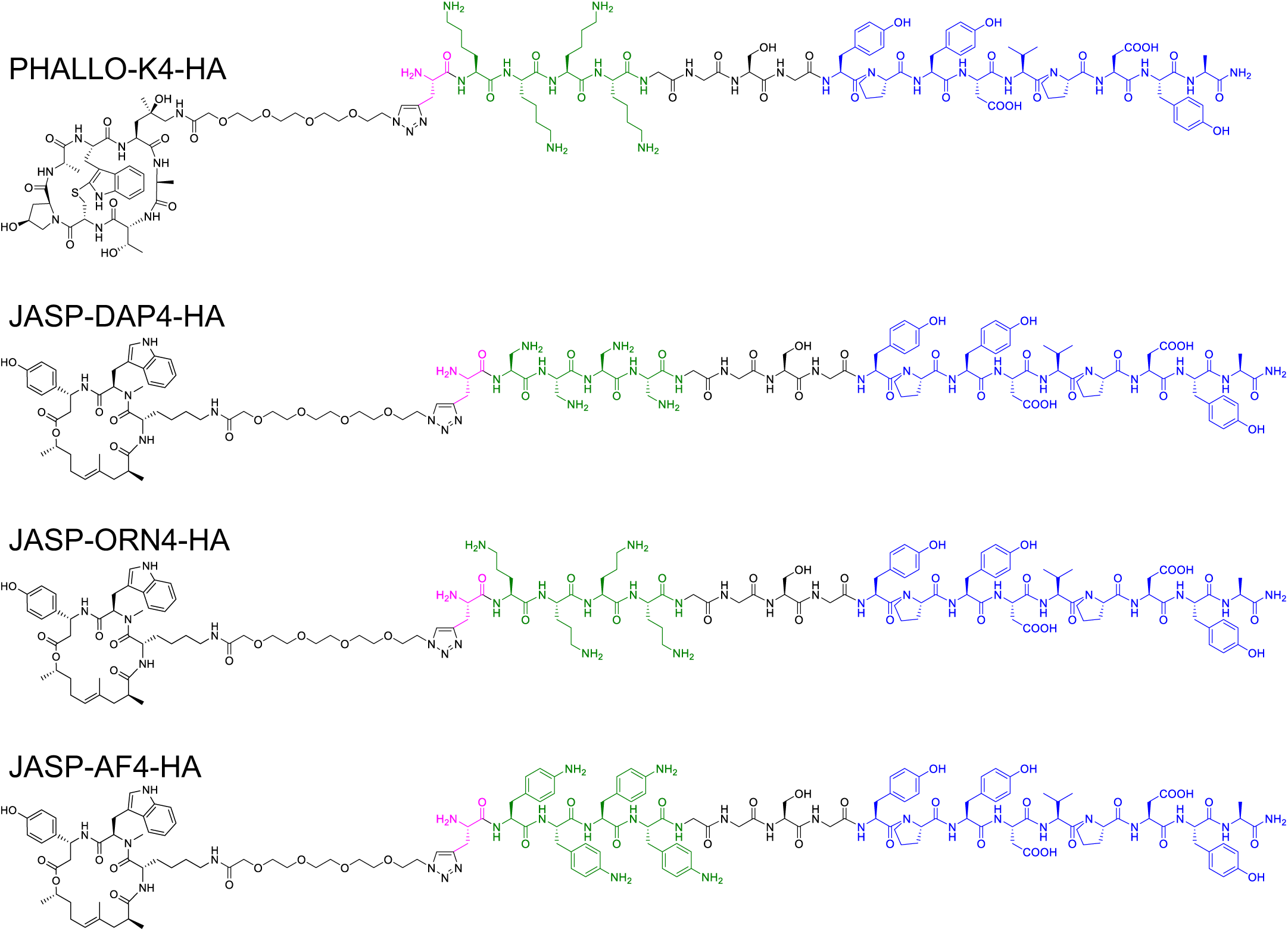
Chemical structure of HAK probes PHALLO-K4-HA, JASP-DAP4-HA, JASP-ORN4-HA and JASP-AF4-HA.

**Supplementary Figure 3.**
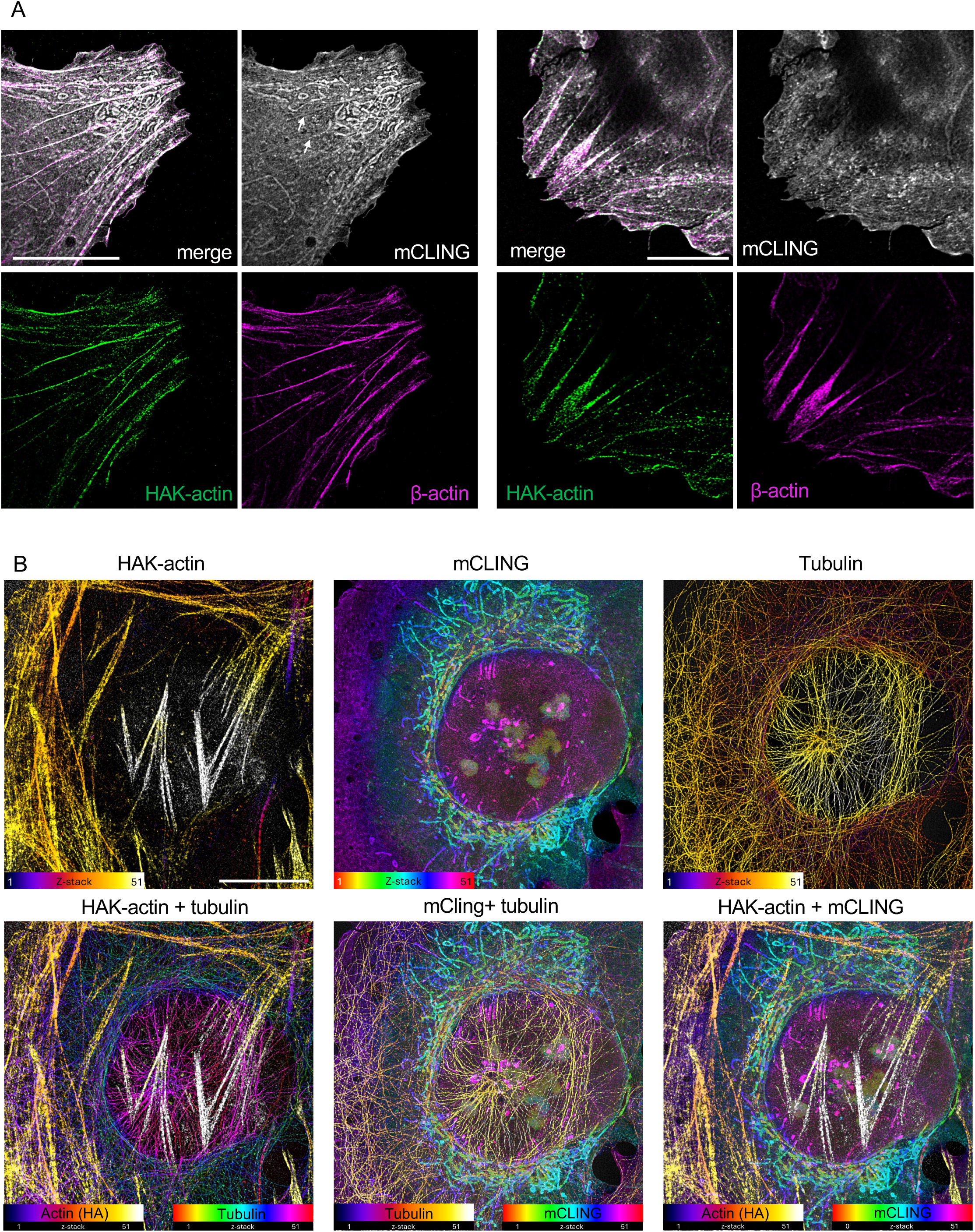
Simultaneous visualization of cellular compartments and cytoskeleton networks. **A**. Confocal images of cryo-fixed and expanded U2OS cells stained with HAK-actin (anti-HA, green), Beta-actin (magenta), and membranes (mCLING, Gray). Scale bars: 10 µm. **B**. Confocal images of cryo-fixed and expanded U2OS stained with HAK-actin (color-coded with “fire” for z positions), mCLING (color-coded with “spectrum” for z positions) and tubulin (color-coded with “fire” or “spectrum” for z positions). All the bottom pictures are a combination of 2 channels. Scale bars: 10 µm

**Supplementary Figure 4.**
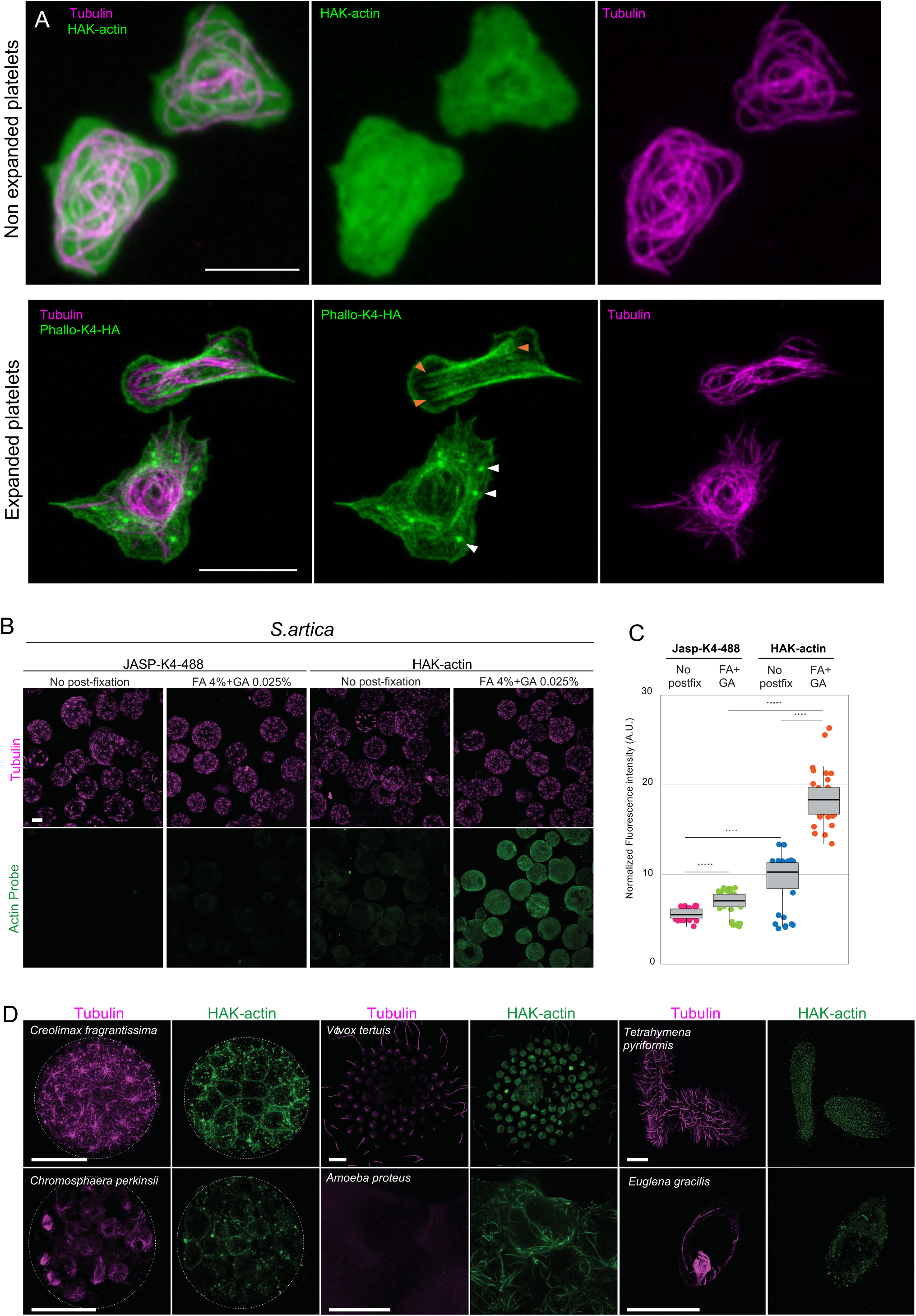
HAK-actin in platelets and microbial eucaryotes. **A**. Widefield image of non-expanded platelets stained for HAK-actin (green) and tubulin (magenta) (top) or expanded platelets stained for PHALLO-K4-HA (green) and tubulin (magenta) (bottom). The split channels images of the expanded platelets shown here correspond to the figure 3E, right. Stress fibers and actin nodules are highlighted with orange and white arrowheads, respectively. Scale bar: 5 µm. **B**. U-ExM samples of *S. arctica* labeled with JASP-K4-488 or HAK-actin probes, either without or with an additional post-fixation step (4%FA + 0.025%GA) prior to the anchoring step (FA/AA). Fluorescence intensity (normalized across images) demonstrates that JASP-K4-488 does not survive the U-ExM protocol and that HAK-actin shows increased signal with the additional crosslinking step. Scale bar: 10 µm. **C**. Box-plot quantifying the fluorescence intensities in (B). N = 35 cells per condition. **D**. U-ExM samples stained with HAK-actin (green) and tubulin (magenta) in six other microbial eukaryotes: the ichthyosporeans *Creolimax fragrantissima* and *Chromosphaera perkinsii*, the green alga *Volvox tertius*, the amoebozoan *Amoeba proteus*, the ciliate *Tetrahymena pyriformis*, and the euglenid *Euglena gracilis*. Note that actin staining is observed in most species except ciliates and euglenids, where phalloidin-stained filamentous actin was never reported. Also note the apparent absence of microtubules in *A. proteus*, which may be associated with a specific cell-cycle stage. Scale bar: 10 µm.

## Data availability statement

The data that support the findings of this study are available from the corresponding authors upon request.

## STAR Methods

### EXPERIMENTAL MODEL

#### Human cell line

*Homo sapiens* bone osteosarcoma U2OS ATCC-HTB-96 were grown in Dulbecco’s modified Eagle’s medium and GlutaMAX, supplemented with 10% fetal calf serum and penicillin and streptomycin (100 μg/ml) at 37 °C in a humidified 5% CO_2_ incubator. All cell cultures were regularly tested for mycoplasma contaminations.

#### Preparation of human platelet rich plasma (PRP)

Whole blood collected in Na-citrate vacutainers (3×3.3 ml/donor) was obtained from the French blood bank. The blood was centrifuged 10 min, 400 g, RT, no break to obtain the upper phase corresponding to the PRP. To increase the platelet concentration of the PRP, 2 ml of the upper phase were removed and kept to prepare plasma (centrifuge 1 min, 12100 g, RT to sediment platelets) and the remaining upper phase was collected as the final PRP.

#### Glutamatergic neuron culture from human iPSCs

Glutamatergic neurons (BrainXell) were seeded at a density of 250 cells/mm² onto coverslips pre-coated overnight at 37°C with poly-D-lysine (0.05mg/mL). The initial culture medium (0 Days In Vitro - DIV) consisted of a basal medium comprising: DMEM/F12 (Sigma) and Neurobasal Medium (Thermo Fisher Scientific), supplemented with N-2 supplement (Thermo Fisher Scientific), K supplement (0.5X, StemCell), SM1 (1X, StemCell), GlutaMax (0.5 mM, Sigma), BDNF (10 ng/mL, PeproTech), GDNF (10 ng/mL, PeproTech), TGF-β1 (1 ng/mL, PeproTech), Geltrex (15 µg/mL, Thermo Fisher Scientific), seeding supplement for Glutamatergic neurons (BrainFast Glut 1X, BrainXell). The BrainPhys medium (Basal medium supplemented with BrainFast D4 (BrainXell)) was gradually introduced on basal medium: 50% at 4DIV, 75% at 7DIV, and 100% at 11DIV. Then the culture media are renewed every two or three days.

#### Mouse retina

All animal experiments were conducted with the authorization numbers GE140 / No national: 34121, according to the guidelines and regulations issued by the Swiss Federal Veterinary Office.

#### Microbial eukaryotes

Among the microbial eukaryotes tested, *Sphaeroforma arctica* (Ichthyosporea), the main species investigated, was cultured as described by^54^. A cryopreserved culture (stored at −80 °C since 2012) was diluted and maintained in Marine Broth 2216 (MB; BD Difco 279110 or Sigma 76448, 37.4 g/L) at 17 °C. Several other microbial eukaryotes were also tested. *Creolimax fragrantissima* (Ichthyosporea) was maintained under similar growth conditions as *S. arctica*. *Chromosphaera perkinsii* (Ichthyosporea) was cultured as previously reported^42^. *Amoeba proteus* (Amoebozoa), *Volvox tertius* (Green algae), *Tetrahymena pyriformis* (Ciliates), and *Euglena gracilis* (Euglenid) were obtained from https://einzeller.izb.unibe.ch/. Upon receipt, these species were immediately chemically fixed in 4% formaldehyde (FA).

### METHOD DETAILS

#### Reagents and antibodies

See Key Resource Table for details regarding reagent and antibodies used in this study.

#### Chemical synthesis

All chemical reagents and anhydrous solvents for synthesis were purchased from commercial suppliers (Sigma-Aldrich, Fluka, Acros, …) and were used without further purification or distillation. The composition of mixed solvents is given by the volume ratio (v/v). ^1^H nuclear magnetic resonance (NMR) spectra were recorded on a Bruker DPX 400 (400 MHz for ^1^H) with chemical shifts (δ) reported in ppm relative to the solvent residual signals (2.50 ppm for DMSO-d6). Coupling constants are reported in Hz. LC-MS was performed on a Shimadzu MS2020 connected to a Nexerra UHPLC system equipped with a Waters ACQUITY UPLC BEH Phenyl 1.7µm 2.1×50mm column. Buffer A: 0.05% HCOOH in H_2_O Buffer B: 0.05% HCOOH in acetonitrile. LC gradient: 10% to 90% B within 6.0 min with 0.5 ml/min flow. Unless otherwise stated, preparative HPLC was performed on a Dionex system equipped with an UltiMate 3000 diode array detector for product visualization on a Waters SymmetryPrep C18 column (7 µm, 7.8 x 300 mm). Buffer A: 0.1% v/v TFA in H2O; Buffer B: acetonitrile. Gradient was from 10% to 90% B within 30 min with 3 ml/min flow.

Peptides used in this study were ordered at Biomatik (www.biomatik.com), with 80-90% purity.

Alkyne-K_4_-HA: H-Pra-(K)_4_GGSGYPYDVPDYA-NH2

Alkyne-DAP_4_-HA: H-Pra-(DAP)_4_GGSGYPYDVPDYA-NH2

Alkyne-ORN_4_-HA: H-Pra-(ORN)_4_GGSGYPYDVPDYA-NH2

Alkyne-AF_4_-HA: H-Pra-(AF)_4_GGSGYPYDVPDYA-NH2

(Pra= propargylglycine, DAP = 1,2-diaminopropionic acid, ORN = Ornithine, AF = 4-aminophenylalanine)

##### NHS-PEG_4_-azide

Azido-PEG_4_-COOH (CAS 201467-81-4) (3 mg, 10.8 µmol, 1.0 eq,) was dissolved in 100 ul DMSO. DIPEA (5 ul, 29.1 µmol, 2.7 eq.) and TSTU (3.9 mg, 13.0 µmol, 1.2 eq.) were successively added and the mixture was incubated at RT for 10 minutes. Reaction completion was monitored by LCMS. The crude reaction mixture was used without further purification and assumed to be a 0.1 M solution of NHS-PEG_4_-azide.

##### Phalloidin-PEG_4_-azide

((R)-4-Hydroxy-4-methyl-Orn⁷)-Phalloidin (CAS 87876-22-0, Bachem) (1.0mg, 1.3 µmol, 1.0 eq.) was dissolved in 100 µl DMSO. DIPEA (2 ul, 11.6 µmol, 8.9 eq.) and crude NHS-PEG_4_-azide (0.1 M in DMSO, 16 ul, 1.6 µmol, 1.2 eq.) were added and the reaction mixture was kept at RT for 15 minutes. The product was purified by preparative HPLC and lyophilized. Yield: 0.9 mg. HRMS (ESI/QTOF) m/z: [M + Na]^+^ Calcd for C_45_H_66_N_12_NaO_15_S^+^ 1069.4384; Found 1069.4332.

##### Jasp-PEG_4_-azide

Boc-Lys-Jasplakinolide derivative [55] (7.8mg, 10 µmol, 1.0 eq.) was dissolved 1 ml TFA/DCM 1:3 and stirred for exactly 10 minutes at RT. 2 ml of toluene were added and the solvents were immediately removed under reduced pressure (water bath at 35°C). The residue was dried under high vacuum for 30 minutes and used without further purification. The residue was dissolved in 300 ul DMSO. DIPEA (20 ul, 116 µmol, 11 eq.) and crude NHS-PEG4-azide (0.1 M in DMSO, 1.2 ml, 12 µmol, 1.2 eq.) were added and the reaction mixture was kept at RT for 15 minutes. The product was purified by preparative HPLC and lyophilized. Yield: 6.4 mg. ^1^H NMR (400 MHz, DMSO) δ 10.80 (d, J = 2.4 Hz, 1H), 9.30 (s, 1H), 8.64 (d, J = 8.8 Hz, 1H), 7.68 (dd, J = 12.5, 8.2 Hz, 2H), 7.59 (t, J = 5.9 Hz, 1H), 7.29 (d, J = 8.1 Hz, 1H), 7.17 – 7.09 (m, 2H), 7.09 – 6.99 (m, 2H), 6.99 – 6.91 (m, 1H), 6.74 – 6.65 (m, 2H), 5.51 (dd, J = 11.3, 5.1 Hz, 1H), 5.18 (ddd, J = 11.5, 8.7, 3.1 Hz, 1H), 4.92 (t, J = 6.6 Hz, 1H), 4.67 (h, J = 6.4 Hz, 1H), 4.60 – 4.52 (m, 1H), 3.86 (s, 2H), 3.59 (dd, J = 5.6, 4.3 Hz, 2H), 3.56 (s, 4H), 3.57 – 3.48 (m, 2H), 3.41 – 3.34 (m, 2H), 3.04 (s, 3H), 2.98 – 2.81 (m, 3H), 2.73 – 2.62 (m, 1H), 2.62 – 2.51 (m, 1H), 2.17 (dd, J = 14.9, 11.3 Hz, 1H), 1.90 – 1.77 (m, 2H), 1.73 (d, J = 14.7 Hz, 1H), 1.56 – 1.43 (m, 1H), 1.49 (s, 3H), 1.43 – 1.30 (m, 1H), 1.16 (d, J = 6.3 Hz, 3H), 1.13 – 1.10 (m, 2H), 0.92 (d, J = 6.7 Hz, 3H), 0.86 – 0.80 (m, 3H). HRMS (ESI/QTOF) m/z: [M + Na]^+^ Calcd for C_48_H_68_N_8_NaO_11_^+^ 955.4900; Found 955.4919.

##### Jasp-PEG_4_-AF488

Jasp-PEG_4_-azide (0.20 mg, 0.30 µmol, 1 eq) was dissolved in 30 µl DMSO, mixed with Alexa488-alkyne (0.25 mg, 0.33 µmol, 1.1 eq.) dissolved in 30 µl DMSO. Separately, 4 µl of TBTA solution (0.1 M in DMSO) was mixed with 4 µl of CuSO_4_ solution (0.1 M in water) and the mixture was added to the alkyne/azide mixture. 8 µl of 0.5 M Sodium ascorbate in water were finally added and the reaction mixture was briefly shaken and incubated under argon for 1h at RT. The product was purified by preparative HPLC and lyophilized. Yield: 0.35 mg, 77%. MS (ESI/single quad) m/z: [M + H_3_]^+3^ Calcd for C_106_H_154_N_21_O_29_S_2_^3+^ 750.0; Found 750.0

##### Alkyne-K_4_-AF488

2-chlorotrityl resin (100 mg, 0.15 mmol, 1 eq.) was swollen in DCM and DMF for 10 min each. The resin was then reacted with a mixture of 2,2′-(Ethylenedioxy)bis(ethylamine) (0.3 ml, 2 mmol, 13.3 eq.), and DMF (1.2 ml) for 30 min at RT. The resin was filtered off and washed with DMF (5 x 3 ml). Standard Fmoc-amino acid SPPS was carried out with this resin to reach the desired sequence Ac-Pra-KKKK-PEG-NH_2_. Briefly, 0.5 mmol Fmoc-Lys(Boc)-OH or Fmoc-propargylglycine-OH, 0.5 mmol of COMU and 0,3 ml DIPEA were dissolved in DMF (1.2 ml). after 5 minutes, the orange-red mixture was added to the resin and incubated for 30 minutes at RT. The resin was filtered off, washed with DMF (5 x 3 ml) and incubated with 20% piperidine in DMF (2.5 ml) for 5 minutes. The resin was filtered off and washed with DMF (5 x 3 ml). After the synthesis, final acetylation step of the N-terminus was carried out by incubating the resin with a mixture of 0.2 ml acetic anhydride, 0.2 ml DIPEA and 1.6 ml DMF for 30 minutes. The resin was filtered off and washed with DMF (5 x 3 ml). The peptide was cleaved from the resin by incubation with hexafluoroisopropanol (1.5 ml) for 10 minutes. The resin was filtered off, washed with DCM (2ml) and the filtrate evaporated. The crude Alkyne-{K(Boc)}_4_-amine peptide (ca 20 mg of crude obtained) was used without further purification for the next step. Crude Alkyne-{K(Boc)}_4_-amine peptide (10 mg, 8.3 µmol, 1eq.) was dissolved in 0.16 ml DMSO, treated with DIPEA (14 ul, 83 µmol, 10 eq.). Separately, AF488 NHS ester (6.1 mg, 8.3 mmol, 1eq.) was dissolved in DMSO (0.16 ml) and added to the peptide-DIPEA solution. The reaction mixture was incubated at r.t. for 3h, purified by preparative HPLC and the solvents were evaporated to dryness. It is to be noted that at this stage, some unlabeled peptide coeluted with the product. The residue was dissolved in TFA/DCM 1:1 (4 ml), stirred for 15 minutes at RT and the solvents were evaporated to dryness. The crude product was used without further purification for the next step. HRMS (ESI/QTOF) m/z: [M + H]^+^ Calcd for C_58_H_85_N_13_O_18_S_2_^+2^ 657.7783; Found 657.7781.

##### Jasp-PEG_4_-K_4_-AF488

Jasp-PEG_4_-azide (1.1 mg, 1.18 µmol, 1 eq) was dissolved in 50 ul DMSO, mixed with Alkyne-K_4_-AF488 (2.2 mg, 1.20 umol, 1 eq.) dissolved in 50 ul DMSO. Separately, 8 ul of TBTA solution (0.1 M in DMSO) was mixed with 8 ul of CuSO_4_ solution (0.1 M in water) and the mixture was added to the alkyne/azide mixture. 16 ul of 0.5 M Sodium ascorbate in water were finally added and the reaction mixture was briefly shaken and incubated under argon for 1h at RT. The product was purified by preparative HPLC and lyophilized. Yield: 1.2 mg. HRMS (ESI/QTOF) m/z: [M + H_2_]^+2^ Calcd for C_106_H_154_N_21_O_29_S_2_^+3^ 749.6882; Found 749.6897.

##### Phalloidin-PEG_4_-K_4_-HA

Phalloidin-PEG_4_-azide (0.45 mg, 0.43 µmol, 1 eq) was dissolved in 50 µl DMSO was mixed with Alkyne-K_4_-HA (1.2 mg, 0.47 µmol, 1.1 eq.) dissolved in 50 µl DMSO. Separately, 4 µl of TBTA solution (0.1 M in DMSO) was mixed with 4 µl of CuSO_4_ solution (0.1 M in water) and the mixture was added to the alkyne/azide mixture. 8 µl of 0.5 M Sodium ascorbate in water were finally added and the reaction mixture was briefly shaken and incubated under argon for 1h at RT. The product was purified by preparative HPLC and lyophilized. Yield: 0.9 mg. HRMS (ESI/QTOF) m/z: [M + H_4_]^+4^ Calcd for C_136_H_205_N_35_O_41_S^+4^ 754.1183; Found 754.1200.

##### Jasp-PEG_4_-K_4_-HA (HAK-actin)

Jasp-PEG_4_-azide (0.42 mg, 0.43 µmol, 1 eq) was dissolved in 40 µl DMSO, mixed with Alkyne-K_4_-HA peptide (1.2 mg, 0.47 µmol, 1.1 eq.) dissolved in 50 µl DMSO. Separately, 4 µl of TBTA solution (0.1 M in DMSO) was mixed with 4 µl of CuSO_4_ solution (0.1 M in water) and the mixture was added to the alkyne/azide mixture. 8 µl of 0.5 M Sodium ascorbate in water were finally added and the reaction mixture was briefly shaken and incubated under argon for 1h at RT. The product was purified by preparative HPLC and lyophilized. Yield: 1.2 mg. HRMS (ESI/QTOF) m/z: [M + H_4_]^+4^ Calcd for C H N O ^+4^ 725.6312; Found 725.6278.

##### Jasp-PEG_4_-DAP_4_-HA

A 10 mM Jasp-PEG_4_-azide in DMSO (100 µl, 1.0 µmol, 1.0 eq) was mixed with a 10 mM Alkyne-Dap_4_-HA peptide solution in DMSO (100 µl, 1.0 µmol, 1.0 eq.). Separately, 5 µl of TBTA solution (0.1 M in DMSO) was mixed with 5 µl of CuSO_4_ solution (0.1 M in water) and the mixture was added to the alkyne/azide mixture. 10 µl of 0.5 M Sodium ascorbate in water were finally added and the reaction mixture was briefly shaken and incubated under argon for 2h at RT. The product was purified by preparative HPLC and lyophilized. Yield: 1.9 mg. HRMS (ESI/QTOF) m/z: [M + H_3_]^+3^ Calcd for C H N O ^+3^ 911.1099; Found 911.1095.

##### Jasp-PEG_4_-ORN_4_-HA

A 10 mM Jasp-PEG_4_-azide in DMSO (100 ul, 1.0 µmol, 1.0 eq) was mixed with a 10 mM Alkyne-Orn_4_-HA peptide solution in DMSO (100 ul, 1.0 µmol, 1.0 eq.). Separately, 5 µl of TBTA solution (0.1 M in DMSO) was mixed with 5 µl of CuSO_4_ solution (0.1 M in water) and the mixture was added to the alkyne/azide mixture. 10 µl of 0.5 M Sodium ascorbate in water were finally added and the reaction mixture was briefly shaken and incubated under argon for 4h at RT. The product was purified by preparative HPLC and lyophilized. Yield: 2.1 mg. HRMS (ESI/QTOF) m/z: [M + H_3_]^+3^ Calcd for C H N O ^+3^ 948.4849; Found 948.4876.

##### Jasp-PEG_4_-4AF_4_-HA

A 10 mM Jasp-PEG_4_-azide in DMSO (100 ul, 1.0 µmol, 1.0 eq) was mixed with a 10 mM Alkyne-4AF_4_-HA peptide solution in DMSO (100 ul, 1.0 µmol, 1.0 eq.). Separately, 5 ul of TBTA solution (0.1 M in DMSO) was mixed with 5 µl of CuSO_4_ solution (0.1 M in water) and the mixture was added to the alkyne/azide mixture. 10 µl of 0.5 M Sodium ascorbate in water were finally added and the reaction mixture was briefly shaken and incubated under argon for 4h at RT. The product was purified by preparative HPLC and lyophilized. Yield: 1.3 mg. HRMS (ESI/QTOF) m/z: [M + H_4_]^+4^ Calcd for C H N O ^+4^ 759.6155; Found 759.6171.

#### Human U2OS cells

##### Actin probes treatment

U2OS cells were grown on 12 mm coverslips. After a quick wash with PBS, they were fixed with 4% paraformaldehyde (PFA) for 5 min at RT and permeabilized with PBS/Tween 0.2% for 10 min at RT. Actin probes were added (100 nM final) after the permeabilization, in PBS/Tween 0.2% for 1h at RT. For few experiments, a post-fixation step consisting in incubation coverslips in 4% PFA + 0.0125% glutaraldehyde (GA) for 15 min at RT was added. After a quick wash with PBS, coverslips were processed for immunofluorescence or U-ExM.

##### Immunofluorescence on non-expanded U2OS cells

After fixation, permeabilization, actin probe treatment, cells were insubated with DAPI (1µg/mL) for 1h RT. Fluoromount was used to mount coverslips on a glass slide.

##### Cryo-fixation of U2OS cells

The cryo-fixation procedure was performed as previously described^3^. Briefly, U2OS cells grown on 12 mm diameter coverslips were plunged freezed into liquid ethane cooled at −170°C using liquid nitrogen. Coverslips were then rapidly transferred into 5ml Eppendorf tubes containing liquid nitrogen pre-cooled acetone. The samples were then incubated overnight on dry ice, with an approximate angle of 45°C, under soft shaking. The samples were subsequently rehydrated by sequential incubations in a mixture of ethanol:water as follows: EtOH 100% (5min, −20°C) - EtOH 100% (5 min, −20°C) - EtOH 95% (3 min, −20°C) - EtOH 95% (3min, −20°C) – EtOH 70% (3min, 4°C) - EtOH 50% (3min, RT) - EtOH 25% (3min, RT) - H2O 100% (RT) - PBS 1x (RT). To ensure cell permeabilization after cryo-fixation, the cells were then incubated with PBS/Tween 0.2% for 10min at RT. Actin probes (100nM final) were either added in acetone before plunging or after the permeabilization, in PBS/Tween 0.2%, for 1h. Then, samples were processed for U-ExM.

##### U-ExM of PFA-fixed or Cryo-fixed U2OS Cells

PFA- or cryo-fixed U2OS cells, treated with actin probes, were then processed for the U-ExM protocol. Briefly, coverslip was first incubated 3h in 1 mL of 2% acrylamide + 1.4% formaldehyde at 37 °C. Then, solution was removed and 35 μL monomer solution composed of 25 μL of sodium acrylate (stock solution at 38% [w/w] diluted with nuclease-free water), 12.5 μL of AA, 2.5 μL of N,N′-methylenebisacrylamide (BIS, 2%), and 5 μL of 10× PBS together with ammonium persulfate (APS) and tetramethylethylenediamine (TEMED) as a final concentration of 0.5% were added on a parafilm. The coverslip was then transferred on the drop of monomer solution (cells facing down) for 5 min at 4 °C followed by 30 min at 37 °C. Then, the gel and the coverslip was detached and immersed in a well of a 6-well plate filled with 2mL of denaturation buffer (200 mM SDS, 200 mM NaCl, 50 mM Tris Base in water (pH 9) for 15 min at RT under shaking to allow the detachment of the gel from the coverslip. The gel was next incubated in 1.5 ml tube filled with denaturation buffer for 1 h 30 at 95 °C. Finally, gel was expanded in three successive ddH2O baths of around 30 min. Round punches of 1 cm diameter were then performed and gels were either frozen in glycerol 50% (in water) or stained after 3 successive baths of PBS. Primary and secondary antibodies were incubated for 2h30 at 37°C in 1,5mL tube in a heat block with shaking in PBS BSA 2%. After each incubation, 3 successive washes with PBS Tween 0,1% of 5 min were done. Stained gels were finally expanded with 3 successive ddH2O baths of 5 min and imaged.

##### iU-ExM of Cryo-fixed U2OS cells

Cryo-fixed U2OS cells were processed for iterative expansion microscopy (iU-ExM) protocol, adapted from established procedures. In brief, cells were fixed in a mixture of 1.4% formaldehyde (FA) and 2% acrylamide (AAm) in 1× PBS for 3 h at 37°C. Gelation was carried out by transferring the coverslip into humidified chamber on ice, where monomer solution containing 10% AAm, 19% sodium acrylate (SA), 0.1% N,N′-dihydroxyethylene bisacrylamide (DHEBA), and 0.25% APS/TEMED was added. After 15 min of incubation on ice, polymerization was completed at 37°C for 45 min. Once gelled, the samples were immersed in denaturation buffer (200 mM SDS, 200 mM NaCl, 50 mM Tris base, pH 6.8) in a 6-well plate until the gel detached from the coverslip. Gels were then transferred to 1.5 mL tubes with fresh denaturation buffer and incubated at 85°C for 1.5 h. This was followed by three 30-min washes in deionized water to initiate expansion. The partially expanded gels were subsequently immunostained and embedded in a second gel for further expansion. For re-embedding, 13 mm punches of the gel were placed in 6-well plates containing 2 mL of a neutral monomer mix (10% AAm, 0.05% DHEBA, 0.05% APS/TEMED) and incubated on ice with gentle agitation for 25 min. Samples were mounted on microscope slides, excess monomer was blotted away using Kimwipes, and a 22 × 22 mm coverslip was applied. Remaining space was filled with monomer solution, and polymerization was carried out in a humidified chamber at 37°C for 1 hour. Following embedding, the gels were treated with an anchoring solution (1.4% FA and 2% AAm) at 37°C for 3 h under shaking, washed with PBS, and then incubated in a second expansion monomer solution (10% AAm, 19% SA, 0.1% BIS, 0.05% APS/TEMED) for 25 min on ice with agitation. Excess solution was removed, the gels were covered with coverslips, and polymerization was completed at 37°C for 1 hour in a humidified environment. To finalize the protocol, the gels were treated with 200 mM NaOH for 1 hour at room temperature with shaking, followed by multiple washes in PBS (∼20 minutes each) until neutral pH was achieved. Full expansion was accomplished through successive incubations in deionized water until the gel size stabilized.

##### Imaging

Image acquisition was performed on an inverted confocal Leica Stellaris 8 microscope or on a Leica Thunder DMi8 microscope using a 20× (0.40 NA) or 63× (1.4 NA) oil objective with Lightning or Thunder SVCC (small volume computational clearing) mode at max resolution, adaptive as “Strategy” and water as “Mounting medium” to generate deconvolved images. 3D stacks were acquired with 0.12 μm z-intervals and an x, y pixel size of 35 nm.

#### Platelet samples

##### Platelet Spreading

Platelet rich plasma (PRP) was diluted in PBS to a concentration of 2.5×10^6^ platelets/ml (plasma concentrations were kept constant at 0.3%) and 400 µl of this suspension was transferred into each well of a 24 well plate containing plasma cleaned coverslips. The plate is centrifuged for 3 min, 600 g at RT to allow synchronized contact of all platelets with the glass surface. The plate is then placed in a cell culture incubator at 37°C. After 60 min platelets are fixed with isotonic formalin (9 volumes formalin / 1 volume 10xPBS) for 15 min at RT and proceed for immunofluorescence and expansion.

##### Platelet Immunolabeling

Fixed platelets were permeabilized with PBS-TritonX100 0.2% for 15 min, washed and incubated in blocking buffer (3% BSA and 10% goat serum in PBS) for 1h at RT. Coverslips were incubated for 60 min with either Jasp-PEG4-HA (1/200) or Phallo-PEG4-HA (1/200) in PBS Tween 0.1%. For non-expanded conditions, samples were washed in PBS and incubated ON at 4°C with mouse-anti tubulin (clone B-5-1-2, 1/500) and rabbit-anti HA (1/500) in blocking buffer. Coverslips were then incubated with secondary antibodies (1/500) for 1h at RT in the dark and mounted on glass slides using Mowiol.

For expanded conditions, samples were washed and incubated 1h30 at RT with mouse-anti tubulin (clone B-5-1-2, 1/250), washed in PBS and PBS/0.2% TritonX100 and incubated with secondary antibody (1/250). Samples were then expanded (see below) and incubated with rabbit-anti HA (1/340) in blocking buffer for 3h 30 min at RT. After 3 washes in PBS PBS/0.2% TritonX100, they were incubated with secondary antibody (1/500) in blocking buffer for 90 min at RT in the dark. Gels were then washed in PBS and PBS/0.2% TritonX100 and expanded 3 times in ddH20.

##### Platelet *Expansion*

After three washing steps, samples were processed for U-ExM essentially as described by ^2,55^. Briefly, stained platelets were incubated in PBS/0.7% formaldehyde/1% acrylamide overnight at RT. The coverslips were then placed upside down onto an ice cold 35 µl drop of gelation solution (PBS/19% Na-acrylate/10% acrylamide/0.1% bisacrylamide) pipetted onto parafilm on ice immediately after addition of TEMED and APS both to a final concentration of 0.5%. After 5 min of incubation on ice, the samples were transferred to 37°C for 1 h. The coverslips plus gel facing up were then incubated in denaturation buffer (5.7% SDS/0.2M NaCI/50mM Tris, pH9) for 15 min at RT with gentle agitation to detach the gel from the coverslip. The gels were then transferred into an eppendorf tube with fresh denaturation buffer and boiled for 30 min at 95°C. Gels were washed in ddH_2_0 twice and incubated in PBS for immunostaining.

##### Platelets data acquisition

Images were acquired using either a wide-field Olympus epifluorescence microscope (BX41) equipped with a Plan 100×/1.25NA oil objective, a DP70 camera and the acquisition software analySIS (non-expanded) or a confocal microscope (Nikon A1R+MP) equipped with a home-made adaptive optics corrector was used with a 40x long distance water immersion objective (U-ExM).

#### Glutamatergic neuron

##### Fixation and actin probe labelling

4DIV and 21DIV glutamatergic neurons were fixed in a mixture of paraformaldehyde and glutaraldehyde (3%-0.1%) in PBS for 20 min at room temperature, washed and incubated in Neurons were incubated in Jasp-PEG4-HA (400 nM) diluted in PBS-Tween20 for 1h at RT.

##### Expansion and immunostaining

Samples were washed and proceeded for expansion as described in^55^. Briefly, Fixed coverslips were incubated in a mixture of acrylamide 2%, Formaldehyde 1.4% diluted in PBS for 3h at 37°C. Gelation was performed on ice by placing the coverslips upside down on a 35mL drop of monomer solution (19% sodium acrylate, 10% acrylamide, 0.1% BIS-acrylamide, 0.5% TEMED and APS). After 5 min incubation on ice, coverslips were placed at 37°C for 30 min in a humid chamber. Gels were next incubated in denaturation buffer for 10 min at RT and transferred in Eppendorf tube at 95°C for 90 min. Gels were washed twice in ddH_2_O and placed in PBS to perform immunostaining. Gels were incubated in PBS-BSA2% in presence of anti-HA, anti-aTubulin and anti-bTubulin (1/250 each) for 4h at RT with agitation. After three washes in PBS-Tween0.1%, gels were incubated in PBS-BSA2% with secondary antibodies anti-rat A594 and anti-GuineaPig A488 for 3h at RT with agitation. Gels were washed in PBS-Tween0.1% and expanded in ddH_2_O.

#### Data acquisition

Images of glutamatergic neurons were acquired using an IX 83 inverted microscope (Olympus), equipped with a Yokagawa CSU-X1 Spinning Disk Unit (Borealis technology) for homogeneous illumination and Ixon3 888 EM-CCD camera (Andor). The oil immersion Plan Apochromat 60x/1.42 NA objective from Olympus was used for all acquisitions.

#### Retina tissue

##### Retina tissue dissection

Mouse eyes were harvested by post-mortem enucleation on adult mice committed to sacrifice. After having performed a small incision in the anterior part, eyes were directly fixed in 4% PFA for 15 min at RT. Then, eyes were immersed in PBS Tween 0.2% for 30 min. Actin probes (200 nM final) were then added in PBS Tween 0,2% for 1h15 at RT. After a quick wash, retinas were then dissected as previously described^32^. Using microscissors, cornea and lens were discarded, and retinas were detached from the sclera. Retinas were then incised on 4 different points to flatten it as a clover inside a 10-mm microwell of a 35-mm petri dish (P35G-1.5-10-C, MatTek) to allow their processing by Ultrastructure expansion microscopy (U-ExM).

##### Retina expansion

Briefly, Retinas were first incubated overnight (ON) in 100 μL of 2% acrylamide + 1.4% formaldehyde at 37 °C. The day after, solution was removed and 35 μL monomer solution (MS) composed of 25 μL of sodium acrylate (stock solution at 38% [w/w] diluted with nuclease-free water), 12.5 μL of AA, 2.5 μL of N,N′-methylenebisacrylamide (BIS, 2%), and 5 μL of 10× PBS was added for 90 min at RT. Then, MS was removed and 90 μL of MS was added together with ammonium persulfate (APS) and tetramethylethylenediamine (TEMED) as a final concentration of 0.5% for 45 min at 4 °C first followed by 3 h incubation at 37 °C. A 24-mm coverslip was added on top to close the chamber. Next, the coverslip was gently removed and 1 ml of denaturation buffer (200 mM SDS, 200 mM NaCl, 50 mM Tris Base in water (pH 9)) was added into the MatTek dish for 15 min at RT with shaking. Then, careful detachment of the gel from the dish with a spatula was performed, and the gel was incubated in 1.5 ml eppendorf tube filled with denaturation buffer for 1 h at 95 °C and then ON at RT. The day after, the gel was cut as a square around the retina that is still visible and expanded in three successive ddH2O baths. Then, the gel was manually sliced with a razorblade to obtain ∼0.5 mm thick transverse sections of the retina that were then processed for immunostaining. For this, primary antibodies were incubated overnight at 4°C, then washed 3 times with PBS Tween 0,1%, and secondary antibodies were incubated 3h at 37°C and also washed 3 times with PBS Tween 0,1%. Finally, gel slices were expanded with 3 baths of 5 min in ddH2O for imaging.

#### Microbial eukaryotes

##### Immunofluorescence on non-expanded S. arctica cells

The cell culture flasks were scraped, and the suspension was transferred to 15 ml Falcon tubes to sediment for 15–30 min. The supernatant was removed, and the cells were transferred to 1.5 ml microfuge tubes, followed by the addition of fixative for 30 min. The cells were fixed with 4% formaldehyde in 250 mM sorbitol solution, washed twice with 1× phosphate-buffered saline (PBS), and resuspended in 20–30 μl of PBS. Permeabilization was performed using six freeze-thaw cycles (liquid N₂, 10 s; 42 °C, 1 min), followed by blocking in 2% bovine serum albumin (BSA) in PBST (1× PBS with 0.1% Tween-20). For actin staining using the actin probes in this study, the probes were diluted 1:500 from a 100 µM stock solution and incubated for 1 h before additional crosslinking with 4% formaldehyde and 0.025% glutaraldehyde for 10 minutes. For antibody labeling, primary antibodies were used at 1:300 and incubated overnight at 37 °C. The antibodies included anti-tubulin rabbit IgG (AA344 and AA345, ABCD Antibodies), anti-actin mouse antibody JLA20 (DSHB), HA-tag mouse monoclonal antibody (12CA5, Invitrogen, MA1-12429), and anti-HA guinea pig IgG (AF291, ABCD Antibodies). This was followed by three washes for 10 min at room temperature and the addition of secondary antibodies. The following secondary antibodies were used at 1:500: Goat anti-mouse IgG (H+L) cross-adsorbed secondary antibody, Alexa Fluor™ 568 (Invitrogen, A-11004); goat anti-guinea pig IgG (H+L) cross-adsorbed secondary antibody, Alexa Fluor™ 488 (Invitrogen, A11073); donkey anti-rabbit IgG (H+L) highly cross-adsorbed secondary antibody, Alexa Fluor™ 488 (Invitrogen, A-21206); donkey anti-rabbit IgG (H+L) highly cross-adsorbed secondary antibody, Alexa Fluor™ 568 (Invitrogen, A10042); and donkey anti-rabbit IgG (H+L) highly cross-adsorbed secondary antibody, Alexa Fluor™ Plus 647 (Invitrogen, A32795). Incubation was performed for 2–5 h at 37 °C. The cells were then washed and resuspended in fresh 1× PBS for imaging. DNA was stained with Hoechst 33352 at a final concentration of 0.4 µM.

##### U-ExM of Microbial Eukaryotes

U-ExM was adapted from Gambarotto et al. and performed as previously described (Shah et al., Nature, 2024; Mikus et al., 2025). Briefly, following fixation, actin-probe incubation, and crosslinking as described previously, the fixed cells were allowed to sediment onto 6 mm poly-L-lysine-coated coverslips for 1 h. This was followed by anchoring in an acrylamide/formaldehyde solution (1% acrylamide, 0.7% formaldehyde) overnight at 37 °C.

A monomer solution (19% (wt/wt) sodium acrylate (Chem Cruz, AKSci 7446-81-3), 10% (wt/wt) acrylamide (Sigma-Aldrich, A4058), 0.1% (wt/wt) N,N′-methylenebisacrylamide (Sigma-Aldrich, M1533) in PBS) was used for gelation, and gels were allowed to polymerize for 1 h at 37 °C in a moist chamber.

For denaturation, gels were transferred to denaturation buffer (50 mM Tris, pH 9.0, 200 mM NaCl, 200 mM SDS) for 15 min at room temperature and then incubated at 95 °C for 1.5 h. Following denaturation, expansion was performed with several water exchanges. After expansion, gel diameter was measured to determine the expansion factor. For all U-ExM images, scale bars indicate actual size, rescaled according to the gel expansion factor.

Immunostaining was performed as described above. All antibodies were prepared in 3% PBS with 0.1% Tween-20. Primary antibodies were incubated overnight at 37 °C, and secondary antibodies were incubated for 2–5 h at 37 °C.

##### Data acquisition

Widefield imaging of *S. arctica* was carried out using a fully motorized Nikon Ti2-E inverted epifluorescence microscope, equipped with a PFS4 hardware autofocus system, a Lumencor SOLA SMII light source, and a Hamamatsu ORCA-spark Digital CMOS camera. Imaging was performed with CFI Plan Fluor objectives: 20× (0.50 NA), 40× (Air), and 60× Oil (0.5–1.25 NA). Confocal microscopy of microbial eukaryotes was conducted using a Leica SP8 upright confocal microscope with an HC PL APO 40×/1.25 glycerol objective.

#### Expansion Factor

Expansion factor (EF) has been calculated by measuring the gel after expansion and dividing the measure by the size of the sample pre-expansion (Range between 4 and 4.3 throughout experiments). All scale bars shown in the paper are corrected to the EF.

### QUANTIFICATION AND STATISTICAL ANALYSIS

#### Quantification

##### Signal Intensity measurement in U2OS

Signal intensity was measured on non-denoised (raw) images with FiJi. For this, a square ROI was drawn by hand in the background (outside of cells) where the mean gray value was measured for both actin probe (488 channel), and Beta-actin (568 channel). Then, the same ROI was used in 3 different locations inside the cell to measure the mean gray value for both actin probe (488 channel), and Beta-actin (568 channel). Finally, the average mean gray value of actin probe signal per cell was divided by the background of actin probe signal on the same image. 5 cells were measured per condition, and 3 independent experiments were performed. To compare actin probe signal with beta-actin signal, the final ratio (signal / background) of actin probe was divided by the ratio (signal / background) of beta-actin.

##### Staining efficiency in S. arctica

Antibody and probe-staining efficiency in *S. arctica* was measured by counting the number of cells with visible actin (488 channel) or tubulin (568 channel) structures. In *S. arctica* actin network can be organized as nodes, filaments or bundles depending on the cell stage, whereas microtubules are present throughout the life cycle. For each condition, between 30 and 246 cells were measured per replicate, and at least three independent experiments, with two technical replicates each, were performed. For anti-actin antibody labelling, only 2 replicates with 2 technical replicates were performed.

##### Signal Intensity measurement in S. arctica

Signal intensity in *S. arctica* was measured from maximum projection images using Fiji. A circular region of interest (ROI) was manually drawn around individual cells, and the mean grey value was quantified for the actin probe (488 channel). A total of 35 cells were measured per condition across two independent experiments.

##### Statistical analysis in S. arctica

All comparisons in Figure 3B and Supplementary figure 5C were performed between two groups based on the figure layout, using a nonparametric Mann-Whitney test. Error bars on the graphs represent standard deviation SD (+/−), and significance levels are indicated as follows: \**P* < 0.05, \*\**P* < 0.01, \*\*\**P* < 0.001, \*\*\*\**P* < 0.0001. All statistical analyses were conducted using R and RStudio.

##### Statistics for U2OS measurements

Comparisons of two groups were realized using the nonparametric Mann-Whitney test. Comparisons of more than two groups were made using the nonparametric Kruskal–Wallis test followed by post hoc test (Dunn’s for multiple comparisons) to identify all the significant group differences. Every measurement was performed on 3 different experiments, with 5 images on each experiment and 3 measurements on each image. Data are all represented as a scatter dot plot with centerline as mean. The graphs with error bars indicate SD (+/−) and the significance level is denoted as usual (\**P* < 0.05, \*\**P* < 0.01, \*\*\**P* < 0.001, \*\*\*\**P* < 0.0001). All the statistical analyses were performed using Prism10.

## ADDITIONAL RESOURCES

### KEY RESOURCES TABLE

| *REAGENT or RESOURCE* | *SOURCE* | *IDENTIFIER* |
| --- | --- | --- |
| *Antibodies* |  |  |
| *Mouse anti-tubulin (clone B-5-I-2)* | *Sigma-Aldrich* | *T5168*<br>*RRID:*<br>*AB_477579* |
| *Guinea Pig anti-αTubulin* | *ABCD antibodies* | *AA344* |
| *Guinea Pig anti-βTubulin* | *ABCD antibodies* | *AA345* |
| *Rabbit Pig anti-αTubulin* | *ABCD antibodies* | *AA344* |
| *Rabbit Pig anti-βTubulin* | *ABCD antibodies* | *AA345* |
| *Rabbit anti-HA (C29F4)* | *Cell Signaling* | *3724*<br>*RRID:*<br>*AB_1549585* |
| *Rat anti-HA (3F10)* | *Roche* | *11867423001*<br>*RRID:*<br>*AB_390918* |
| *Mouse anti-HA Monoclonal (12CA5)* | *Invitrogen* | *#MA1-12429* |
| *Guinea Pig anti-HA* | *ABCD antibodies* | *AF291* |
| *Alexa Fluor 488 Goat anti-guinea pig IgG* | *Invitrogen* | *A11073* |
| *Alexa Fluor 594 Goat anti-rat IgG* | *Invitrogen* | *A11007* |
| *Alexa Fluor 488 goat anti-mouse IgG* | *Invitrogen* | *A11029* |
| *Cy3 goat anti-rabbit IgG* | *Jackson* | *111-165-003* |
| *Mouse beta-Actin* | *Proteintech* | *#60008-1-Ig* |
| *Mouse beta-Actin* | *Proteintech* | *#66009-1-Ig* |
| *Mouse beta-Actin* | *DSHB* | *#JLA20* |
| *Goat anti-Mouse IgG (H+L) Cross-Adsorbed Secondary Antibody, Alexa Fluor™ 568* | *invitrogen* | *Catalog #A-11004* |
| *Donkey anti-Rat IgG (H+L) Highly Cross-Adsorbed Secondary Antibody, Alexa Fluor™ 488* | *Invitrogen* | *Catalog #A-21208* |
| *Goat anti-guinea pig IgG (H+L) Cross-Adsorbed Secondary Antibody, Alexa Fluor™ 488* | *Invitrogen* | *Catalog #A-11073* |
| *Donkey anti-Rabbit IgG (H+L)*<br>*Highly Cross-Adsorbed*<br>*Secondary Antibody, Alexa*<br>*Fluor™ 488* | *Invitrogen* | *Catalog #A-*<br>*21206* |
| *Donkey anti-Rabbit IgG (H+L)*<br>*Highly Cross-Adsorbed*<br>*Secondary Antibody, Alexa*<br>*Fluor™ 568* | *Invitrogen* | *Catalog #A-*<br>*10042* |
| *Donkey anti-Rabbit IgG (H+L)*<br>*Highly Cross-Adsorbed*<br>*Secondary Antibody, Alexa*<br>*Fluor™ Plus 647* | *Invitrogen* | *Catalog #A-*<br>*32795* |
| *Biological samples* |  |  |
| *Platelet rich plasma* | *French Blood Bank* |  |
| *Chemicals, peptides, and recombinant proteins* |  |  |
| *Formalin* | *Sigma-Aldrich* | *HT5012* |
| *Formaldehyde (FA)* | *Sigma-Aldrich* | *F8775-25ML* |
| *Paraformaldehyde 16% (PFA)* | *EMS* | *15710* |
| *Glutaraldehyde (GA)* | *Sigma-Aldrich* | *G5882* |
| *Phallo-4K-HA* | *This study* | *This study* |
| *JASP-4K-HA* | *This study* | *This study* |
| *Triton-X100* | *MERK* | *11869* |
| *Tween-20* | *Sigma-Aldrich* | *P1379* |
| *BSA* | *Euromedex* | *1005-7* |
| *Goat Serum* | *Thermo Fisher Scientific* | *16210-064* |
| *Acrylamide (AA)* | *Sigma-Aldrich* | *A4058-100ML* |
| *N,N'- methylenbisacrylamide 2% (BIS)* | *Sigma-Aldrich* | *M1533* |
| *Sodium Acrylate 97-99% (SA)* | *Sigma-Aldrich or homemade<sup>26</sup> or AK Scientific* | *408220 R624* |
| *Ammonium Persulfate (APS)* | *Thermo Fisher Scientific* | *17874* |
| *Tetramethylethyldiamine (Temed)* | *Thermo Fisher Scientific* | *17919* |
| *Poly-D-Lysine (U-ExM)* | *Sigma-Aldrich* | *P7280 / P1024* |
| *Poly-D-Lysine (culture cell)* | *Sigma-Aldrich* | *P6407* |
| *DMEM/F12* | *Sigma-Aldrich* | *D8437* |
| *Neurobasal Medium* | *Thermo Fisher Scientific* | *21103049* |
| *N-2 supplement* | *Thermo Fisher Scientific* | *17502048* |
| *B-27 Supplement* | *Thermo Fisher Scientific* | *17504044* |
| *K supplement* | *BrainXell* | *BX-2020* |
| *SM1* | *StemCell* | *#05711* |
| *GlutaMax* | *Sigma-Aldrich* | *G8541* |
| *BDNF* | *PeproTech* | *450-02* |
| *GDNF* | *PeproTech* | *450-10* |
| *TGF-β1* | *PeproTech* | *100-21C* |
| *Geltrex* | *Thermo Fisher Scientific* | *A1413202* |
| *BrainFast Glut* | *BrainXell* | *BX-2300* |
| *BrainFast D4* | *BrainXell* | *BX-2040* |
| *Acetone, 99.8%, Extra Dry* | *AcroSeal™, Thermo Scientific Chemicals* | *326801000* |
| *mCLING* | *Synaptic Systems* | *#710 006AT1* |
| *Deposited data* |  |  |
| *Experimental models: Cell lines* |  |  |
| *Cortical glutamatergic neurons* | *BrainXCell* | *BX-0300* |
| *U2OS* | *ATCC* | HTB-96 |
| *Experimental models: Organisms/strains* |  |  |
| *S. arctica* | *Dudin Lab; ATCC* | *ATCC; PRA-297* |
| *C. fragrantissima* | *Dudin Lab; ATCC* | *ATCC; PRA-284* |
| *C. perkinsii* | *Dudin Lab* | *42,56* |
| *Amoeba proteus* | https://einzeller.izb.unibe.ch/ | *101* |
| *Volvox tertius* | https://einzeller.izb.unibe.ch/ | *112* |
| *Tetrahymena pyriformis* | https://einzeller.izb.unibe.ch/ | *104* |
| *Euglena gracilis* | https://einzeller.izb.unibe.ch/ | *111* |
| *Software and algorithms* |  |  |
| *Fiji* | https://imagej.net/software/fiji/downloads |  |
| *GraphPad Prism 10* | https://www.graphpad.com |  |
| *R Studio* | https://rstudio-education.github.io/hopr/starting.html |  |
| *Other* |  |  |
| *ethane:propane 37%:63%* | *PanGas* |  |

